# SRF transcriptionally regulates the oligodendrocyte cytoskeleton during CNS myelination

**DOI:** 10.1101/2022.09.21.508765

**Authors:** Tal Iram, Miguel A. Garcia, Jérémy Amand, Achint Kaur, Manasi Iyer, Mable Lam, Nicolas Ambiel, Andreas Keller, Tony Wyss-Coray, Fabian Kern, J. Bradley Zuchero

## Abstract

Myelination of neuronal axons is essential for nervous system development. Myelination requires dramatic cytoskeletal dynamics in oligodendrocytes, but how actin is regulated during myelination is poorly understood. We recently identified serum response factor (SRF)—a transcription factor known to regulate expression of actin and actin regulators in other cell types—as a critical driver of myelination in the aged brain. Yet, a major gap remains in understanding the fundamental role of SRF in oligodendrocyte lineage cells. Here we show that SRF is required cell autonomously in oligodendrocytes for myelination during development. Combining ChIP-seq with RNA-seq identifies SRF-target genes in OPCs and oligodendrocytes that include actin and other key cytoskeletal genes. Accordingly, SRF knockout oligodendrocytes exhibit dramatically reduced actin filament levels early in differentiation, consistent with its role in actin-dependent myelin sheath initiation. Together, our findings identify SRF as a transcriptional regulator that controls the expression of cytoskeletal genes required in oligodendrocytes for myelination. This study identifies a novel pathway regulating oligodendrocyte biology with high relevance to brain development, aging, and disease.

**Highlights:**

- Developmental CNS myelination requires the transcription factor SRF in oligodendrocytes.
- SRF targets actin and actin-regulatory but not myelin related genes.
- SRF drives oligodendrocyte actin cytoskeleton dynamics during early stages of myelination.

## INTRODUCTION

Myelination is essential for rapid nerve signaling and has emerged as an important regulator of central nervous system (CNS) development, plasticity, and disease (Monje, 2018). The cellular mechanisms controlling myelin formation and dynamics are still incompletely understood. Oligodendrocytes form myelin by dramatically rearranging their cell morphology to ensheath and then spirally wrap dozens of individual myelin sheaths around neuronal axons. Central to the ability of an oligodendrocyte to form myelin is its actin cytoskeleton (Brown and Macklin, 2020), which powers morphological changes in two distinct steps: first, actin assembly is required for early stages of myelination in which oligodendrocytes extend their cellular processes to make first contact with axons that they loosely ensheath (Wilson and Brophy, 1989; Zuchero et al., 2015) similar to how actin assembly drives the extension of neuronal growth cones (Fox et al., 2006). Second, and unexpectedly, dramatic disassembly of the oligodendrocyte actin cytoskeleton occurs prior to the start of myelin wrapping (Nawaz et al., 2015; Zuchero et al., 2015). The first stage—ensheathment—requires expression of proteins that promote actin filament assembly, including Arp2/3 and its regulators. In contrast, the subsequent stage—wrapping—does not require actin assembly factors, while actin *disassembly* factors (e.g. cofilin, ADF, gelsolin) are all highly upregulated and are required for wrapping (Nawaz et al., 2015; Zuchero et al., 2015). Thus, precise control over actin assembly and disassembly—e.g. through tight regulation of gene expression of actin regulatory proteins—is likely to be essential to coordinate oligodendrocyte morphology changes required for these subsequent steps of myelination. The mechanisms regulating the oligodendrocyte cytoskeleton during myelination remain largely unknown.

Serum Response Factor (SRF) is a MADS box transcription factor that functions as a major transcriptional regulator of the actin cytoskeleton in diverse cell types (Miano et al., 2007) including neurons (Knoll and Nordheim, 2009), muscle (Esnault et al., 2014), and cardiac cells (Deshpande et al., 2022; Hauschka, 2001). Many of the cytoskeletal genes that are induced during oligodendrocyte differentiation (Cahoy et al., 2008; Dugas et al., 2012; Zhang et al., 2014) are known targets of SRF (Sun et al., 2006). In the developing nervous system, SRF is required in neural precursor cells for specification of oligodendrocyte precursor cells (OPCs) and astrocytes (Lu and Ramanan, 2012). In addition, neuronal SRF was found to affect myelination non-cell autonomously by controlling neuronal secretion of paracrine signals that affect oligodendrocyte differentiation (Stritt et al., 2009). However, whether SRF also plays a direct, cell-autonomous role in oligodendrocytes during myelination is unknown.

We recently discovered that infusing cerebrospinal fluid (CSF) from young mice into the brains of old mice improves memory function by a mechanism that appears to be largely dependent on the formation of new myelin (Iram et al., 2022). Young CSF increased OPC proliferation, differentiation into oligodendrocytes, and the number of myelinated axons in the hippocampus. Mechanistically, we found that the major cellular target of young CSF in oligodendrocytes is SRF. SRF expression is rapidly induced in OPCs treated with young CSF, followed within hours by the induction of numerous known SRF target genes. Additionally, young CSF increased actin filament levels and growth cones in OPCs, consistent with activating SRF. In culture, the young CSF-induced increase in OPC proliferation is dependent on SRF, as SRF-KO OPCs failed to proliferate in response to young CSF. However, it is still unknown whether SRF also regulates myelination itself—for example by controlling expression of actin genes known to be required in oligodendrocytes for ensheathment and/or wrapping of axons.

Here, we show that SRF is required cell-autonomously in oligodendrocytes for the formation of myelin during development. Mechanistically, the direct gene targets of SRF in OPCs and oligodendrocytes include actin itself and several genes that regulate the formation and stability of actin filaments. Loss of SRF in cultured oligodendrocytes causes a dramatic reduction in actin filament levels early in differentiation, a time point in which actin filament assembly is required for axon ensheathment. Together, our results reveal mechanistic insight into how the oligodendrocyte cytoskeleton is regulated to promote myelination during development and provide a novel pathway that could be targeted to boost, protect, and restore myelination in aging or disease.

## RESULTS

### SRF is expressed by OPCs and oligodendrocytes during CNS development

We first sought to determine when in the oligodendrocyte lineage SRF is expressed during CNS development. We used multiplexed fluorescence RNA in situ hybridization (RNAScope) to visualize and quantify expression of SRF transcripts in brain sections from postnatal day 8 (P8) and P16 mice (Figure 1A), time points representing the start of myelination in the mouse brain. Simultaneous labeling of PDGFRa (OPCs) and Olig2 (all oligodendrocyte lineage) allowed us to quantify SRF expression in both OPCs (Olig2-expressing, PDGFRa-high) and differentiating or mature oligodendrocytes (Olig2-expressing, PDGFRa-low) while avoiding other cell types that express SRF (Figures 1B and 1C). Consistent with published transcriptomics studies (Marques et al., 2016; Zhang et al., 2014) and our prior work on SRF in the aging brain (Iram et al., 2022), SRF mRNA was expressed by both OPCs and oligodendrocytes throughout the brain at both P8 and P16 (Figure 1D). Furthermore, immunostaining with a knockout-validated antibody confirmed SRF protein expression in oligodendrocyte nuclei (Figure 1E), consistent with its known function as a transcription factor. We confirmed SRF’s nuclear localization by immunostaining primary cultured OPCs and oligodendrocytes using the same antibody (Figure 1F). Thus, SRF is expressed by both OPCs and oligodendrocytes during postnatal development, positioning it to regulate gene expression during myelination.

**Figure 1.**
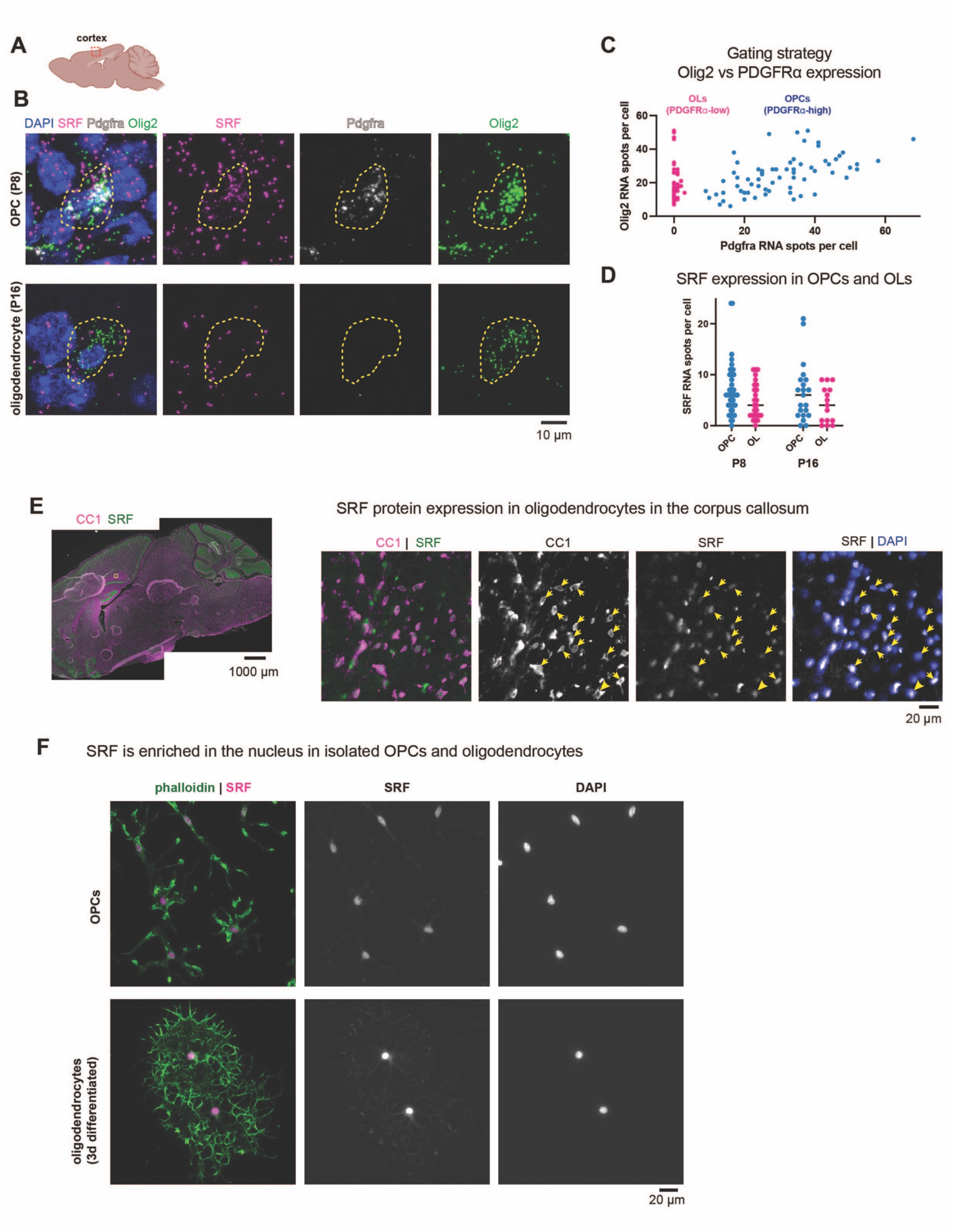
Expression of SRF in the oligodendrocyte lineage. A. Region of interest in sagittal sections of P8 and P16 brains. B. RNAscope for Pdgfra, Olig2 and SRF in the cortex of P8 and P16 brains. n=2, scale bar represents 10μm. C. Gating strategy for selection of OPCs (Pdgfra-high) and oligodendrocytes (OLs; Pdgfra-low) expressing cells. D. Quantification of SRF spots per cell in OPCs and oligodendrocyte population at P8 and P16. Each dot represents a cell, bar represents the mean from all cells. n=3 mice at each age. E. Overview of immunostaining of a P16 mouse with CC1 (mature oligodendrocytes) and SRF and an enlargement of the corpus callosum. Scale bar represents 1000μm and 20μm in the enlargement. F. SRF and phalloidin staining in isolated OPCs or oligodendrocytes (OL) differentiated for 3 days in culture. Scale bar represents 20μm.

### Conditional knockout of SRF causes CNS hypomyelination

To test the role of SRF in myelination in vivo, we generated mice in which a floxed allele of SRF (Ramanan et al., 2005) was conditionally deleted from OPCs using Olig2-Cre (Lappe-Siefke et al., 2003) (Figures S1A). SRF-fl/fl; Olig2-Cre/+ conditional knockout mice (hereafter “SRF-cKO”) were born in Mendelian frequencies and displayed no gross behavioral or motor defects. We confirmed complete loss of SRF mRNA in OPCs using RT-PCR on purified OPCs from SRF-cKO mice and littermate controls (“SRF-flox,” genotype: SRF-^flox/flox^; and “SRF-cHet,” genotype: SRF-^flox/+^; Olig2-^Cre/+^; Figure S1B). In vivo, immunostaining revealed loss of SRF protein from oligodendrocytes, but not from neighboring cells (e.g. neurons) (Figure S1C).

Having confirmed loss of SRF from oligodendrocyte cells in SRF-cKO mice, we used transmission electron microscopy to test whether SRF is required in oligodendrocytes for myelination. We focused on the optic nerve, a uniformly-myelinated axon tract that we and others have studied extensively as a model of CNS myelination (Dangata and Kaufman, 1997; Nawaz et al., 2015; Snaidero et al., 2014; Zuchero et al., 2015). At P18 (during active myelination), SRF-cKO nerves had on average half as many myelinated axons as control SRF-flox littermates with no other abnormalities (Figures 2A and 2B). This degree of hypomyelination of SRF-cKO mice persisted into adulthood (Figures 2D and 2E), indicating that the reduction of myelinated axons in cKOs was not just due to a transient developmental delay. Focusing on only myelinated axons, morphometric analysis revealed that myelin thickness and *g*-ratio were both unaffected in SRF-cKO mice (Figures 2C, 2F, and S2A-S2D) although SRF-cKO myelin was slightly biased towards large-caliber axons—suggesting a specific defect in ensheathing smaller caliber axons (Figures S2E-S2H). Thus, while SRF is important for axonal ensheathment, it appears to be dispensable for the later stage of myelin wrapping.

**Figure 2.**
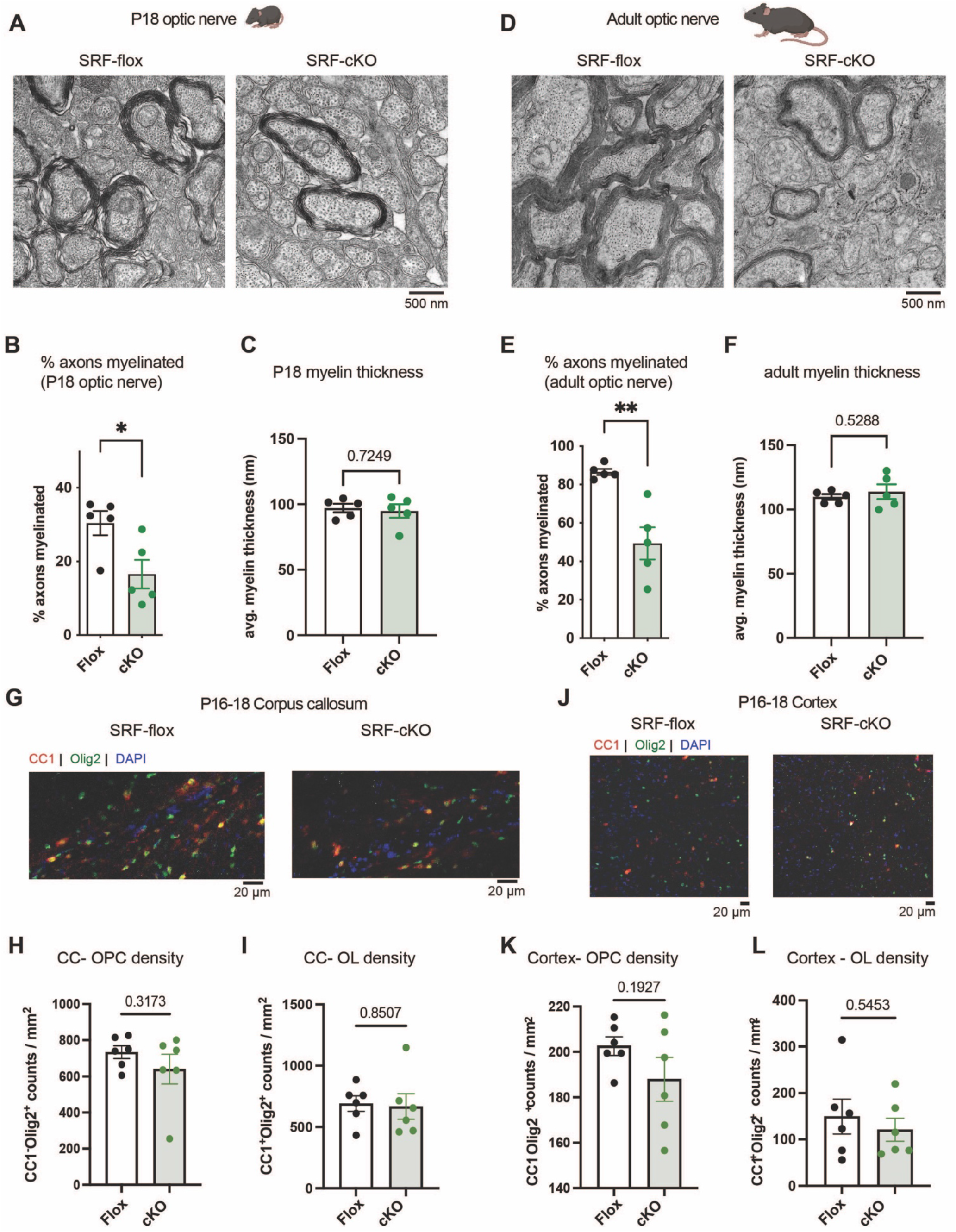
SRF is required in oligodendrocytes for developmental myelination. A. Transmission electron microscopy of optic nerves from p18 SRF-flox and SRF-cKO. Scale bar represents 500nm. B. Quantification of precent axons myelinated at P18. Each datapoint is an average of at least 700 axons per nerve from 5-6 micrographs spanning the entire nerve. n=5 animals from each genotype. Unpaired t-test; * p=0.0262. C. Quantification of myelin thickness at P18. Each datapoint is an average of at least 700 axons per nerve from 5-6 micrographs spanning the entire nerve. n=5 animals from each genotype. Unpaired t-test. D. Transmission electron microscopy of optic nerves from adult SRF-flox and SRF-cKO mice. Scale bar represents 500nm. E. Quantification of precent axons myelinated in adult mice. Each datapoint is an average of at least 700 axons per nerve from 5-6 micrographs spanning the entire nerve. n=5 animals from each genotype. Unpaired t-test; ** p = 0.0024. F. Quantification of myelin thickness in adult mice. Each datapoint is an average of at least 700 axons per nerve from 5-6 micrographs spanning the entire nerve. n=5 animals from each genotype. Unpaired t-test. G. Representative images of CC1 (red) and Olig2 (green) staining of corpus callosum (CC) of P16-P18 SRF-flox and SRF-cKO mice. Scale bar represents 20μm. H. Quantification of OPC density (CC1^-^Olig2^+^ / mm^2^) in corpus callosum (CC) of P16-P18 SRF-flox and SRF-cKO mice. n=6 animals from each genotype. Unpaired t-test. I. Quantification of oligodendrocyte density (CC1^+^Olig2^+^ / mm^2^) in corpus callosum (CC) of P16-P18 SRF-flox and SRF-cKO mice. n=6 animals from each genotype. Unpaired t-test. J. Representative images of CC1 (red) and Olig2 (green) staining of cortex of P16-P18 SRF- flox and SRF-cKO mice. scale bar represents 20μm. K. Quantification of OPC density (CC1^-^Olig2^+^ / mm^2^) in cortex of P16-P18 SRF-flox and SRF-cKO mice. n=6 animals from each genotype. Student T-Test. L. Quantification of oligodendrocyte density (CC1^+^Olig2^+^ / mm^2^) in cortex of P16-P18 SRF-flox and SRF-cKO mice. n=6 animals from each genotype. Unpaired t-test.

The hypomyelination phenotype could be a result of a defect in differentiation or viability of mature oligodendrocytes. We therefore quantified the cell density of OPCs (CC1^-^Olig2^+^ cells) and oligodendrocytes (CC1^+^Olig2^+^ cells) in the corpus callosum (CC, Figures 2G-2I) and cortex (Figures 2J-2L). We found similar densities of OPCs and oligodendrocytes in both regions with a slight trend towards fewer OPCs in the cortex (Figure 2K), suggesting that SRF is not necessary for oligodendrocyte maturation and cell viability. This was in line with similar levels of MBP staining intensity in the cortex and corpus callosum (Figure S2-I-2K). Together, these results indicate that the hypomyelination phenotype in SRF-cKO mice is a result of improper myelination and that SRF is required cell-autonomously within oligodendrocytes for the initial steps of myelination.

### SRK-cKO OPC and oligodendrocyte RNAseq identifies significant alterations in cytoskeletal genes

To gain better mechanistic understanding of the role of SRF in myelination, we performed transcriptome analysis of primary OPCs purified from SRF^flox/flox^ mice, induced with a Cre-expressing virus *ex vivo* and sequenced at the proliferative stage (OPC) or on differentiation day 3 (immature oligodendrocytes). We identified 157 differentially expressed genes (p.adj<0.05) in SRF-KO OPCs vs SRF-WT and 264 in SRF-KO oligodendrocytes, including Srf which was significantly downregulated in both datasets (Figures 3A, 3C, and Supplementary tables 1-2). Gene set enrichment analysis (GSEA) of SRF-KO OPC genes identified depletion of pathways associated with “Transcription regulation” and “Cell cycle”, in accordance with the role of SRF as a transcriptional regulator driving cell proliferation (Iram et al., 2022). Notably, enriched pathways contained genes like Apoe and Clusterin (Clu) associated with diseases such as “Alzheimer’s Disease” and “Parkinson’s Disease” (Figure 3B). SRF-KO oligodendrocytes depleted pathways included “actin cytoskeleton”, “postsynaptic density” and “myelin sheath” (Figure 3D). Lastly, we plotted known SRF targets (by the TRANSFAC database) (Fu and Weng, 2004) and found downregulation of genes associated with the actin cytoskeleton, including upstream regulators such as Rhoj and actin genes such as Actb (Figures 3E-3H). Thus, SRF broadly regulates gene expression in OPCs and oligodendrocytes. We next sought to determine which of these genes are directly regulated by SRF.

**Figure 3.**
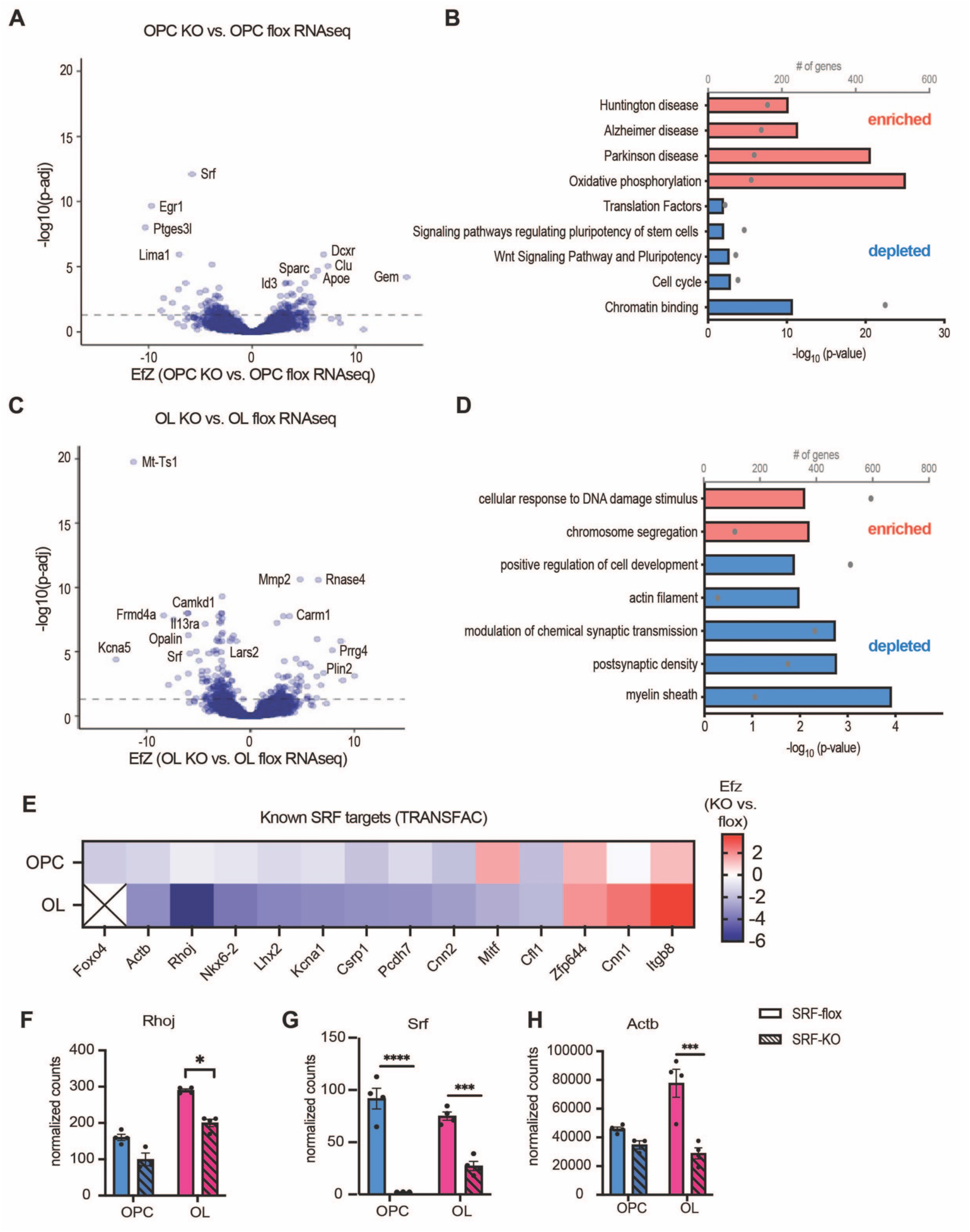
Functional role of SRF in the oligodendrocyte lineage. A. Volcano plot of differentially expressed genes of SRF-KO OPCs vs. SRF-flox OPCs. The dashed line represents Padj=0.05. SRF-flox n=4, SRF-KO n=3. B. Pathways enriched (red) and depleted (blue) in SRF-KO OPCs. C. Volcano plot of differentially expressed genes of SRF-KO oligodendrocytes (OL) vs. SRF-flox OLs. The dashed line represents Padj=0.05. n=4. D. Pathways enriched (red) and depleted (blue) in SRF-KO oligodendrocytes. E. Heatmap of selected SRF targets (predicted by the TRANSFAC database) showing the effect size of KO vs. flox OPCs and oligodendrocytes. F-H. Normalized expression counts of Rhoj (F), Srf (G) and Actb (H) in SRF-flox and SRF-KO OPCs and oligodendrocytes. OPC SRF-KO n=3. All the rest n=4. Statistics by Deseq2.

### ChIP-seq identifies SRF target genes in oligodendrocytes

Based on the finding that SRF target genes were transcriptionally modulated in SRF-KO OPCs and oligodendrocytes, we used chromatin immunoprecipitation-sequencing (ChIP-seq) to identify direct SRF gene targets. Due to the high number of cells required to perform transcription factor ChIP-seq, we performed the experiments on primary rat OPC cultures under proliferation conditions (OPC) or on day 3 of differentiation (immature oligodendrocytes). We identified 848 significant SRF binding sites (peaks) in total and narrowed down the list to 329 high-confidence targets that appeared consistently across replicates within the same group and not in the IgG control (Figure 4A and supplementary table 3). Consistent with other SRF ChIP-seq datasets (Esnault et al., 2014), most peaks were enriched within 2kb of the transcription start site (TSS) (Figure 4B). Inspection of the target sequences confirmed an enrichment for the SRF CArG consensus (Figure 4C) as well as for Thap11 and Zic2 motifs (Supplementary table 3). Roughly half (n=60) of the target genes with a confirmed CArG consensus were shared between OPCs and oligodendrocytes and included known SRF targets such as immediate early genes (Egr1, Egr2, Egr3, Fos, Junb and Srf) and cytoskeleton genes (Actb, Actng1, Arc, Vcl) (Figure 4D). Among the OPC unique target genes we found several genes with promyelinating effects such as Cardiotrophin-1 (Ctf1) (Stankoff et al., 2002), and involved in nervous system development such as Reticulon 4 (Rtn4/Nogo) and UTP11. Notably, among the oligodendrocyte unique target genes we found cytoskeletal proteins such as Filamin A (Flna) and Pppr12b and members of the postsynaptic density scaffold Homer1 and Dlg4. Next, we plotted the ChIP-seq CArG genes that were significantly altered in SRF-KO OPCs and oligodendrocytes (pval< 0.05, Figure 4E). Actin genes such as beta actin (Actb) were downregulated in SRF-KO OPCs and oligodendrocytes indicating that they are indeed directly regulated primarily by SRF and therefore downregulated when SRF is knocked-out. Interestingly, we did not identify “myelin genes” as direct SRF targets. Together, these experiments identified genes controlled by SRF in OPCs and oligodendrocytes and highlighted actin cytoskeletal regulation as a potential role of SRF during myelination.

**Figure 4.**
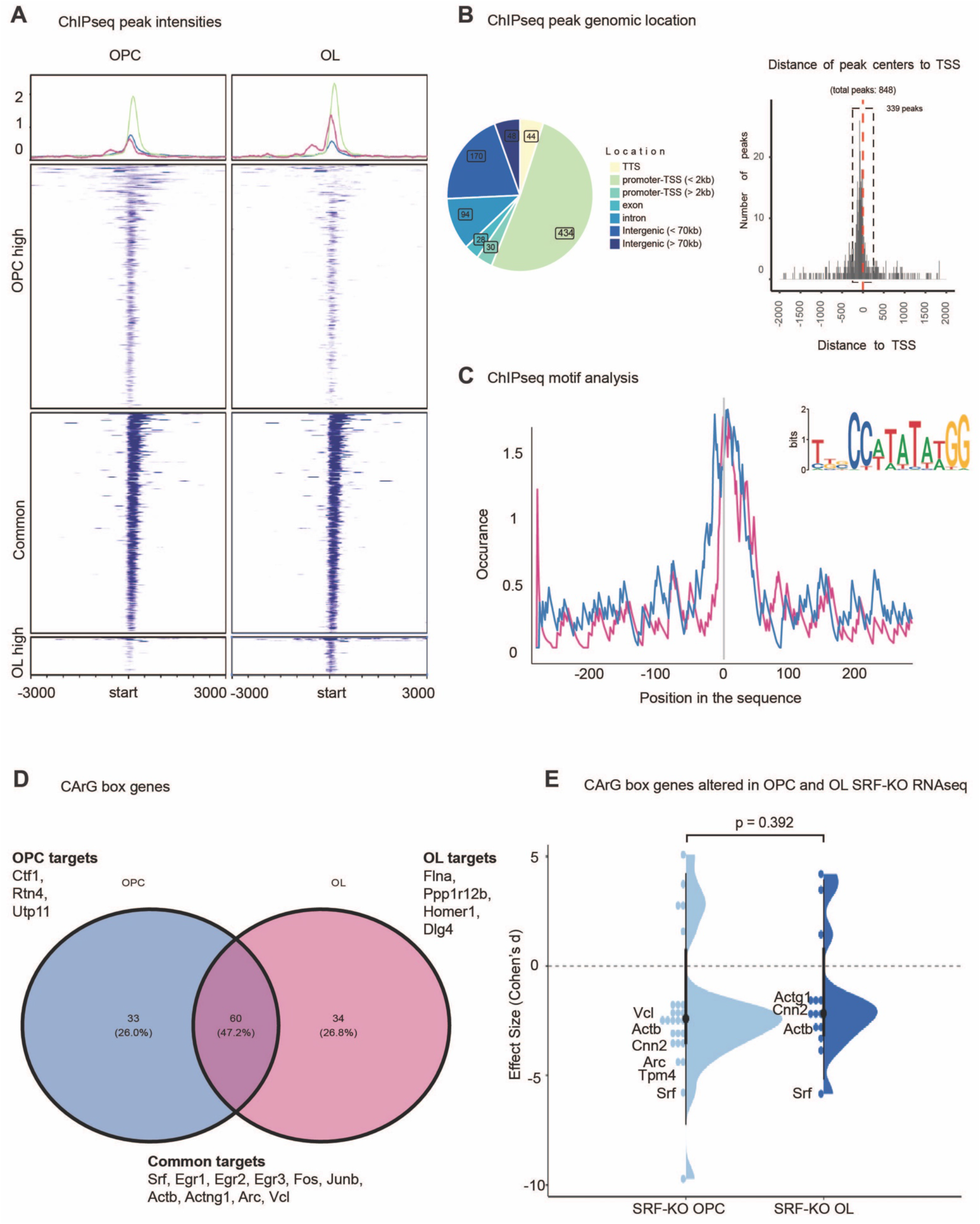
ChIP-seq identifies SRF target genes in OPCs and oligodendrocytes. A. Heatmap of SRF ChIP-seq peak intensities in OPC and oligodendrocyte samples. OPCs n=4, oligodendrocytes n=3. B. ChIP-seq peak genomic location of peaks showing that most of the peaks localize to the promoter and transcription start site (TSS). C. ChIP-seq motif analysis identifying CarG box, the SRF binding motif in OPC and oligodendrocyte peaks. D. Venn-diagram of genes associated with the CarG box peaks in OPC and oligodendrocyte samples. E. Violin plot of CarG box genes in SRF-KO vs. SRF-flox OPC and oligodendrocyte RNASeq. Wilcoxon rank-sum test.

### SRF regulates the oligodendrocyte actin cytoskeleton

Actin dynamics are critical for CNS myelination, with the early (SRF-dependent) stage of ensheathment requiring actin filament assembly (Nawaz et al., 2015; Zuchero et al., 2015). SRF is known in other cell types to regulate actin filament assembly and disassembly by transcriptional control of numerous actin regulatory genes (Esnault et al., 2014; Miralles et al., 2003; Sotiropoulos et al., 1999). Our ChIP-seq and RNA-seq results suggested that, in oligodendrocytes, SRF primarily regulates genes that promote actin assembly (including actin itself) and not genes that regulate actin disassembly (Supplementary Figure S4). To test this, we purified OPCs from SRF-WT and SRF-cKO littermates and compared their actin cytoskeletons over a time course of differentiation (Figure 5A). In OPCs and early stages of oligodendrocyte differentiation, SRF-cKO cells had dramatically reduced actin filament levels (~20-40% of SRF-WT levels; Figures 5B-5E). However, both SRF-WT and SRF-cKO oligodendrocytes underwent a similar degree of actin disassembly during late stages of differentiation, and by full maturation at 7 days of differentiation cells from both genotypes were nearly indistinguishable (Figure 5D). Furthermore, expression of the differentiation marker and major myelin protein MBP was not different between SRF-WT and SRF-cKO oligodendrocytes. These data suggest that the effect of SRF on the actin cytoskeleton is due to its direct regulation of actin gene expression rather than a more general effect on oligodendrocyte differentiation.

**Figure 5.**
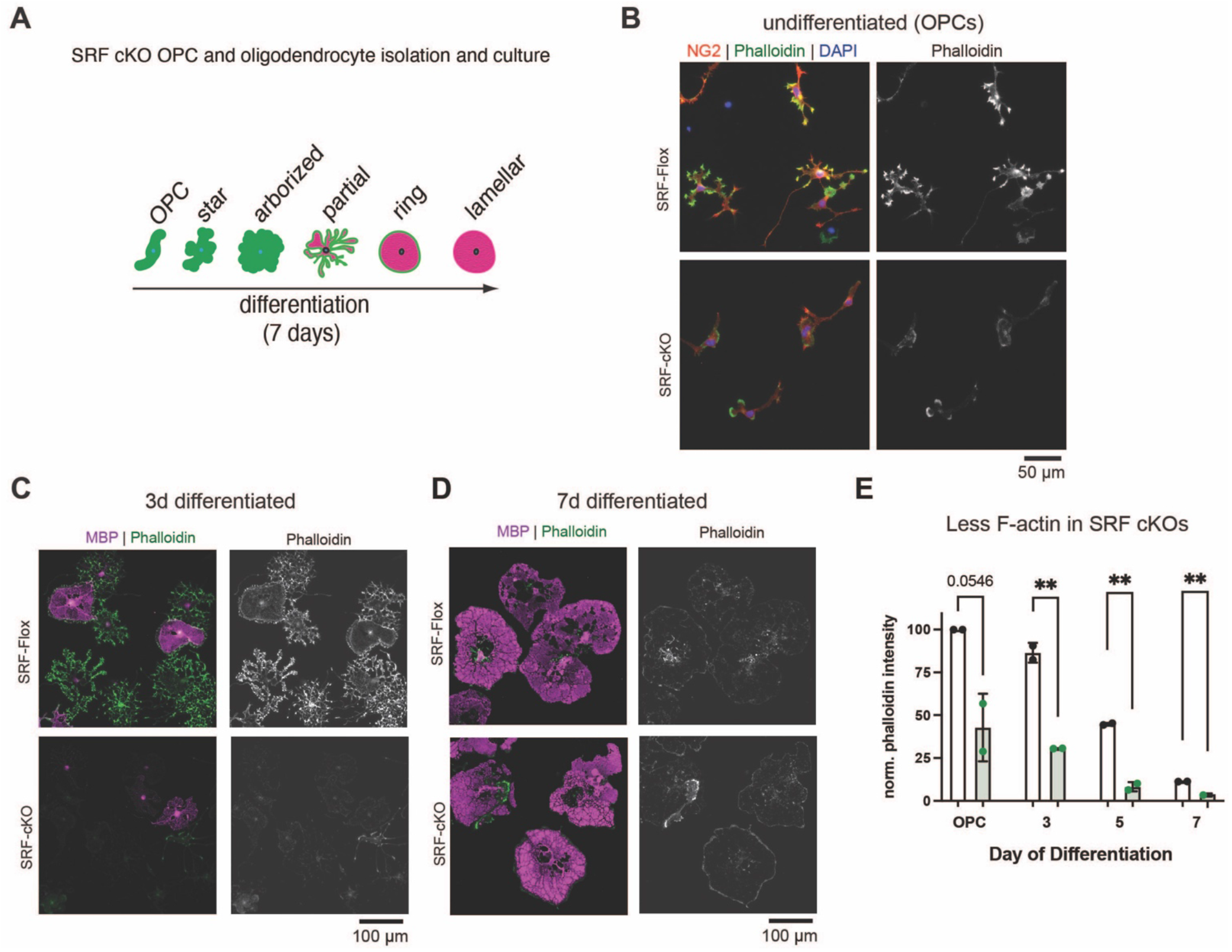
SRF regulates the oligodendrocyte cytoskeleton. A. Diagram of OPC to oligodendrocyte morphology changes when differentiated in culture. B. SRF-flox and SRF-cKO OPCs stained for the OPC marker NG2 (red), phalloidin (green) and DAPI (blue). Scale bar represents 50μm. C. SRF-flox and SRF-cKO immature oligodendrocytes (3 day differentiated) stained for MBP (magenta), phalloidin (green) and DAPI (blue). Scale bar represents 100μm. D. SRF-flox and SRF-cKO mature oligodendrocytes (7 day differentiated) stained for MBP (magenta), phalloidin (green) and DAPI (blue). Scale bar represents 100μm. E. Quantification of phalloidin intensity in SRF-flox and SRF-cKO at different stages of differentiation. n=2 preps. Unpaired t-test; ** p<0.01.

Together, our results indicate that SRF promotes early stages of developmental myelination—at least in part—by controlling the expression of genes required for actin filament assembly.

## DISCUSSION

Myelin plays essential roles in CNS development, dynamics, and disease, but the mechanisms that regulate its formation are still incompletely understood. In particular, actin cytoskeleton dynamics precisely regulate the morphological changes oligodendrocytes must undergo to build myelin sheaths, but the molecules that regulate actin in oligodendrocytes are largely unknown. Here, we show that the transcription factor SRF is required in oligodendrocytes for early stages of myelination. SRF is expressed developmentally by both OPCs and oligodendrocytes throughout the CNS. SRF-cKO mice have severe defects in numbers of axons myelinated, consistent with an ensheathment defect, while myelin sheaths that do form wrap normally. Combining ChIP-seq with RNA-seq of SRF knockout oligodendrocytes identifies SRF target genes during myelination, and include genes required for the formation of actin filaments. Accordingly, SRF knockout oligodendrocytes fail to assemble actin normally, potentially explaining their inability to ensheath axons. Thus, SRF is a transcription factor that promotes myelination by regulating the expression of actin and actin-regulatory genes during the early, actin-dependent stage of myelination.

SRF is a versatile regulator of a multitude of cellular functions in various tissues like skeletal muscle (Braun and Gautel, 2011), heart (Guo et al., 2018) and brain (Knoll and Nordheim, 2009). In the developing brain, SRF modulates hippocampal neuronal growth cone integrity, migration, axon guidance and circuit assembly (Knoll et al., 2006). Mechanistically, at least half of all known SRF gene targets encode proteins that regulate actin-dependent processes in mammalian cells (Medjkane et al., 2009). It is interesting to speculate that SRF regulates neuronal and oligodendrocyte development by modulating their cytoskeleton to promote each cell’s unique structure and function.

How does SRF regulate myelination? Studying Arp2/3 cKO mice, we previously formulated a two-step model of myelination in which actin assembly drives oligodendrocyte process outgrowth and ensheathment at the start of myelination, while at later stages actin disassembly drives myelin wrapping (Zuchero et al., 2015). In the current study, SRF-cKO mice show very similar phenotypes to ArpC3 cKO mice, including hypomyelination in the optic nerve (Figure 2) and reduced actin filament levels in oligodendrocytes (Figure 5). These similarities, along with the ChIP-seq and RNAseq of SRF-KO OPCs and oligodendrocytes, suggest that a major role of SRF is to regulate the expression of target genes required for actin filament assembly at the actin-dependent stage of early myelination. Of note, Knoll and colleagues hypothesized that SRF could also drive myelination directly in oligodendrocytes by binding to myelin gene promoters (Stritt et al., 2009). This was supported by another study which showed that pharmacologic inhibition of SRF inhibited OPC differentiation (Buller et al., 2012). However, our ChIP-seq studies found no evidence that SRF directly regulates the expression of classical myelin genes, but rather promotes actin formation which is required for proper axon ensheathment (Figure 4). It remains possible that loss of SRF causes a subtle downstream delay in oligodendrocyte differentiation indirectly—e.g. via other transcription factors that it regulates, or even as a consequence of reduced actin levels.

Interestingly, in other cell types, SRF’s transcriptional activity is directly mediated by cofactors that respond to the balance of actin monomer/filament in the cytoplasm (Connelly et al., 2010; Knoll, 2010; Vartiainen et al., 2007). In this way, it has been proposed that SRF could act as an “actin homeostat” to respond to the state of the actin cytoskeleton by inducing gene expression of proteins that, in turn, regulate actin assembly or disassembly. Although the actin-dependent regulation of SRF has—thus far— been only observed in cultured cells, it is tempting to speculate that such a regulatory feedback circuit could function in cells to coordinate cytoskeletal changes with gene expression. An ideal cell type to study actin-SRF feedback may be the oligodendrocyte, a cell type that undergoes dramatic cytoskeleton rearrangements during its differentiation from actin-rich to nearly devoid of actin filaments (Nawaz et al., 2015; Zuchero et al., 2015). In future studies, it will be interesting to explore whether SRF responds to cytoskeletal changes during oligodendrocyte differentiation to coordinate gene expression with cytoskeletal remodeling.

Beyond development, myelin loss and the progressive inability of oligodendrocytes to regenerate myelin are increasingly found to underly cognitive defects associated with aging and neurodegenerative disease. We have recently found that SRF is downregulated in aged mouse OPCs, and that infusion of young CSF promotes SRF pathway activation and actin assembly in OPCs, as well as OPC proliferation and differentiation (Iram et al., 2022). In the current study, we gained more mechanistic insight into the functional outcomes of SRF loss. We found that SRF-cKO in oligodendrocytes leads to developmental hypomyelination which persisted to adulthood. Future studies focusing on adult induced SRF-KO in OPCs and oligodendrocytes could unravel the important roles of SRF in adaptive myelination in the context of learning and memory and cognitive aging. Furthermore, in our SRF-KO RNAseq dataset, expectedly, SRF and many SRF targets such as Egr1 were transcriptionally downregulated. However, to our surprise, some disease related genes such as Apoe and clusterin were upregulated (Figure 3A-B). Interestingly, a recent preprint identified a subset of clusterin expressing OPCs in the 5xFAD mouse model of Alzheimer’s Disease. They further described that clusterin acts as a potent inhibitor of OPC differentiation and expression of myelin proteins (Beiter et al., 2022). Combined, these results may link SRF loss to a dysfunctional OPC disease state that appears in aging and neurodegenerative diseases.

In this study, we gained a deeper understanding of the roles of SRF in regulating oligodendrocyte maturation during development. Ultimately, a full mechanistic understanding of how oligodendrocytes undergo such dramatic morphology changes to form myelin will open the door to understanding how myelin is dynamically remodeled during learning and could reveal novel approaches for regenerating myelin in aging and disease.

## METHODS

### Animals and ethics statement

All procedures involving animals were approved by the Institutional Administrative Panel on Laboratory Animal Care (APLAC) of Stanford University and followed the National Institutes of Health guidelines. Mice were group-housed under standard 12:12 light-dark cycles at temperatures of 18-23 degrees Celsius and 40-60% humidity with free access to food and water, and disposable bedding in plastic cages. All mice received regular monitoring from veterinary and animal care staff and were not involved in prior procedures or testing. Sprague-Dawley rats and C57BL/6 mice were ordered from Charles River Laboratories. SRF-flox mice (Ramanan et al., 2005) (Jax strain #: 006658) were a kind gift of Prof. David Ginty (Harvard University) and Olig2-Cre mice (Schuller et al., 2008) were a kind gift of Prof. David Rowitch (University of Cambridge). All mouse lines were maintained by breeding with C57BL/6 mice. Both male and female mice were studied for all in vivo experiments. For cell culture studies with OPCs from mutant mice, brains of both sexes were pooled to obtain sufficient cell numbers.

### RNA fluorescence in-situ hybridization (RNAscope)

C57Bl6/J (WT) mice were euthanized by decapitation with (P16) or without (P8) isoflurane anesthesia to allow for rapid collection of tissues for RNAscope. The brain was removed from the skull and immediately frozen on dry ice. Tissues were kept at −80 °C until sectioning. For tissue sectioning, frozen brains were mounted in O.C.T. compound (Thermo Fisher Scientific, 23-730-571) until sectioning. 10–12 μm thick sagittal sections were collected with a Leica CM3050 S cryostat preset to −20 °C. Sections were immediately transferred to cold Superfrost Plus (VWR, 48311-703) microscopy slide and stored at −80 °C until staining assays were performed.

RNAscope multiplex fluorescence (ACDbio) assays were completed according to the manufacturer’s sample preparation manuals. Multiplex fluorescence kits (ACDbio, 320851) and the HybEZTM II Hybridization system (ACDbio, 321710) were used in all RNAscope assays. Fresh frozen tissue slices were fixed for 15 minutes in 4% paraformaldehyde solution (EMS, 15710) and serially dehydrated in ethanol washes immediately before assay.

After dehydration steps, samples were dried at room temperature for 20 minutes and then treated with pretreat IV (ACDbio, 320842) for 30 minutes. RNAscope probes were hybridized to target mRNAs, and the samples were treated with hybridization reagents Amp-FL 1-4 and counterstained with DAPI. Samples were covered with microscopy cover glass (Fisherbrand, 12-544-E) and mounted with Prolong Gold antifade mount (Invitrogen, 10144).

### RNAscope Probe List

RNAscope Probe List:
| Probe ID | Species | ACDbio Cat# |
| --- | --- | --- |
| positive control (Polr2a, PPIB, UBC) | mouse | 320881 |
| negative control (Dapb) | Bacillus subtilis | 320871 |
| Pdgfra | mouse | 480661 |
| Olig2 | mouse | 447091 |
| SRF | mouse | 574761 |

For RNAscope imaging and analysis, slides were imaged with a Zeiss LSM 800 laser scanning confocal microscope using Zeiss Zen Blue software. A 63X oil objective was used to obtain Z-stack images of tissues. Specimens were blinded in the experiments and unblinded after imaging and analysis. Cells were chosen randomly for imaging based on DAPI or Olig2 expression without viewing the SRF channel. Quantification of the culture or in vivo RNAscope experiments was performed automatically using FishQuant (Mueller et al., 2013). Except for cases in which a significant background signal was noted in a slide, the same threshold analysis settings were applied to all specimens within an experiment. Cell boundaries were determined using DAPI staining and mRNA localization.

### Immunostaining of mouse tissue

Antibodies used in this study include rabbit-anti-SRF H-300 (Santa Cruz sc13029; 1:100), mouse-anti-CC1 (Millipore, OP980; 1:100), Rat-anti-MBP (Abcam, ab7349; 1:100; knockout-validated (Zuchero et al., 2015), rabbit-anti-NG2 (Millipore AB5320; 1:250), Rabbit-anti-Olig2 (Millipore AB9610, 1:200) and highly cross-absorbed Alexa Fluor 488-, 594-, or 647-labeled secondary antibodies (Thermo Fisher).

Tissues on slides were incubated with 10% goat serum (Thermo Scientific, PCN5000) in 0.1% Triton (Sigma, T8787) containing PBS for 1 hour. Primary antibody incubation was done in 1% goat serum in PBS overnight, then slides were rinsed 4 times for 30 minutes with 0.1% Triton containing PBS. Secondary antibody solution (Alexa Fluor 488, 1:500) was done at room temperature for 1-2 hours. Slides were rinsed four times with 0.1% Triton containing PBS for 30 minutes. Slides were counterstained with DAPI and mounted with Prolong Gold antifade mount (Invitrogen, 10144).

### Transmission electron microscopy sample preparation

Transmission electron microscopy (TEM) was completed in the Stanford Cell Sciences Imaging Facility. Samples were prepared according to previously published protocols (81). Samples were initially washed in chilled Karlsson-Schultz fixative (2.5% glutaraldehyde, 4% PFA in phosphate buffer, pH 7.3) and incubated in 2% OsO4 for four hours at 4 °C. Samples were then serially dehydrated at 4 °C and embedded in EmBed812 (EMS, 14120). 80 nm sections were taken using an UC7 (Leica, Wetzlar, Germany) and were collected on formvar/Carbon coated 100 mesh Cu grids. Sections were then stained for 40 seconds in 3.5% uranyl acetate in 50% acetone followed by staining in Sato’s lead citrate for 2 minutes. TEM images were obtained using a JEOL JEM-1400 120kV with a Gatan OneView 4k X 4k digital camera. Quantification of TEM images was performed manually using Fiji/Image-J, blinded to genotype.

### Purification and culture of oligodendrocytes

Primary oligodendrocyte precursor cells (OPCs) were purified by immunopanning from P5-P7 Sprague Dawley rat or P6-P7 transgenic mouse brains as previously described (Dugas et al., 2012; Emery and Dugas, 2013). OPCs were typically seeded at a density of 150,000-250,000 cells/10-cm dish and allowed to recover and proliferate for 4 days in culture before lifting cells via trypsinization and distributing for proliferation or differentiation assays. All plasticware for culturing oligodendrocyte precursors were coated with 0.01 mg/ml poly-D-lysine hydrobromide (PDL, Sigma P6407) resuspended in water. All glass coverslips for culturing oligodendrocyte precursors were coated with 0.01 mg/ml PDL, which was first resuspended at 100x in 150 mM boric acid pH 8.4 before diluting to 1x in water (PDL-borate). To proliferate primary oligodendrocyte precursors, cells were cultured in serum-free defined media (DMEM-SATO base medium) supplemented with 4.2 μg/ml forskolin (Sigma-Aldrich, Cat#F6886), 10 ng/ml PDGF (Peprotech, Cat#100-13A), 10 ng/ml CNTF (Peprotech, Cat#450-02), and 1 ng/ml neurotrophin-3 (NT-3; Peprotech, Cat#450-03) at 37°C with 10% CO2. To induce differentiation, cells were switched to DMEM-SATO base media containing 4.2 μg/ml forskolin (Sigma-Aldrich, Cat#F6886), 10 ng/ml CNTF (Peprotech, Cat#450-02), 40 ng/ml thyroid hormone (T3; Sigma-Aldrich, Cat#T6397) and (for mouse OPCs only) 1x NS21-MAX (R&D Systems AR008).

### Confirming SRF knockout in OPCs/oligodendrocytes

We used the following primers to genotype SRF knockout mice: SRF-forward-2: TGCTGGTTTGGCATCAACT and SRF-reverse: GGCACTGTCTCAGGGTGTCT (WT SRF results in a 400 bp band; SRF-floxed allele results in a 650 bp band); Cre-forward: GCTAAGTGCCTTCTCTACACCTGC and Cre-reverse: GGAAAATGCTTCTGTCCGTTTG (presence of Cre indicated by the presence of a 500 bp band). For RT-PCR to analyze gene expression and test SRF knockout in OPCs, RNA was purified from immunopanned (95-99% pure) and proliferated OPC samples prepared as above, using the QIAGEN RNeasy Mini Kit. Note that SRF is expressed in cell types other than OPCs, so we used purified OPCs rather than total brain lysate for these experiments. Equal amounts of RNA (50-400 ng total) were reverse transcribed using SuperScriptIII (Invitrogen). Equal volumes from RT samples to be compared were then amplified using Platinum Taq (Invitrogen) with 26 PCR cycles, and sub-plateau reactions were analyzed by densitometry (NIH ImageJ). Primer sequences were SRF-RT-for: ACCAGTGTCTGCTAGTGTCAGC and SRF-RT-rev: CATGGGGACTAGGGTACATCAT.

### Immunofluorescence of primary rat or mouse oligodendrocytes

Primary rat OPCs (WT) or primary mouse OPCs from SRF-flox and SRF-cKO littermate mice were harvested and proliferated for 1-4 days as described above in “Purification and Culturing of Cells”. Cells were seeded onto 12-mm glass coverslips (Carolina Biological Supply No. 63-3029) coated with PDL-borate (see above) at a density of 10,000 cells/coverslip in differentiation media. At the specified day of differentiation, cell media was removed and coverslips were fixed with 4% formaldehyde in PBS for 15 minutes exactly, gently washed 3x in PBS, permeabilized with 0.1% Triton-X-100 in PBS for 3 minutes exactly, and gently washed 3x in PBS to remove Triton, all at room temperature (RT). Permeabilized cells were then blocked with 3% BSA (Sigma-Aldrich, A2153) in PBS for 20 minutes at RT. Antibody conditions for staining cultured oligodendrocytes were identical to those listed above in “Immunostaining of mouse tissue.” Primary antibody incubation was performed at 4°C overnight in 3% BSA/PBS. Coverslips were incubated with secondary antibodies (1:1000 in 3% BSA/PBS) for 2 hours at RT. Actin filaments were stained using Alexa Fluor-conjugated phalloidin (Invitrogen; 50 nM in PBS) for 15 min exactly at RT. Cells were gently rinsed three times in PBS between each staining step. Finally, coverslips were mounted with Fluoromount G with DAPI (SouthernBioTech, 0100-20) on Superfrost microscopy slides (Fisherbrand, 12-550-143), air-dried overnight in the dark at RT, then stored at −20°C until shortly before imaging. Cells were imaged by widefield epifluorescence with a Zeiss Axio Imager M1 and Axiovision software using a 20x 0.8 NA Plan Apo objective (Carl Zeiss Microscopy). Images were acquired blinded to the genotype/experimental condition with identical illumination and acquisition conditions per biological replicate. Images were analyzed through batch processing in Fiji/Image J (Schindelin et al., 2012). Cells from 2-4 biological replicates were analyzed (2 coverslips per N served as technical replicates). To quantify actin filament content in cells, images were first color-thresholded to create cell-based ROIs. Then, mean cellular phalloidin intensity was measured, and background intensity of a region of the same image with no cells was subtracted from this intensity value.

### ChIP-seq of OPCs and oligodendrocytes

Rat OPCs were allowed to proliferate for 6 days to reach 40 million cells in multiple 15cm plates. Half of the plates were fixed at the OPC stage while the rest were switched to differentiation medium and fixed on day 3 of differentiation. Replicates in this experiment were independent OPC preps. Chromatin immunoprecipitation was performed following iDeal ChIP-seq kit for Transcription Factors (Diagenode, Cat. No. C01010055) using 7 cycles for chromatin shearing on a Bioruptor Pico sonicator (Diagenode, Cat. No. B01060001). Prior to sonication, each sample was split to 300ul reactions (containing each roughly 4 million cells) in 1.5 ml Bioruptor Microtubes (Cat. No. C30010016), sonicated, spun 16,000g for 10 min and 270ul of supernatant was collected to a new tube and stored in −80 deg. One aliquot of each sample was used for immunoprecipitation with 5ul of Rabbit-anti-SRF antibody that was validated for ChIP (SRF (D71A9) XP® Rabbit mAb, #5147) and an additional OPC and oligodendrocyte sample from each group for immunoprecipitation with an IgG control antibody (provided in the kit).

Libraries for next-generation sequencing were prepared using MicroPlex Library Preparation Kit v3 x48 rxns (C05010001, Diagenode) using 12 PCR cycles on a 96-plate thermal cycler (Biorad) and purified without size selection.

Library quantity and quality was assessed using a Bioanalyzer (Agilent) and Qubit. Libraries were pooled and sequenced on a Nextseq550 sequencer (Illumina) using single end 75bp for Read 1 and 8bp for index 1 and 8bp for Index 2 with a high output 75bp kit (20024906, Illumina).

In pilot studies, Chromatin Shearing Optimization kit - Low SDS (iDeal Kit for TFs) (Cat. No.C01020013) was used to determine optimal number of sonication shearing cycles to yield 200-300 bp fragments.

### Bioinformatics processing of ChIP-seq data

ChIP-seq data analysis was performed using the *nextflow-core ChIP-seq v1.2.1* pipeline (https://doi.org/10.5281/ZENODO.3966161) with default parameters, unless otherwise stated. In short, the pipeline first performs adapter trimming to the raw single-end reads using *Trim Galore* (option --nextseq=20) (https://www.bioinformatics.babraham.ac.uk/projects/trim_galore/). Next, reads were mapped to the *Rnor 6.0* rat genome obtained from Illumina iGenomes (Ensembl) using *BWA (Li and Durbin, 2009). MACS2* (Zhang et al., 2008) was configured to call narrow peaks with a Benjamini-Hochberg FDR of 0.05 and an effective genome size of 2 × 10^!^. *HOMER (Heinz et al., 2010)* was then used to annotate peaks with information from the closest known genomic feature based on the Ensembl database. Consensus peaks for OPC and oligodendrocytes were aggregated by considering peaks present in at least one replicate of the respective cell type. We performed known and *de novo* motif detection in the −250 to +250 nt regions around detected peaks using *MEME-ChIP* (*Bailey et al., 2015*). Next, FIMO (Bailey et al., 2015) was used to get individual motif matches for MEME-ChIP results. The matrix of peak intensity around transcription start sites (TSS) was calculated using *deepTools (Ramirez et al., 2016).* Data transformation and plotting were performed using R v4.1.3.

### RNA-seq of OPCs and oligodendrocytes

Mouse OPCs were purified from brains of SRF floxed by immunopanning as described above for rat OPCs. On day 3 of the culture, SRF-f/f OPCs were split and plated in 12-well plates at a density of 30K per well. When the cells were in suspension in proliferation media before plating, 1*10^10^ viral genomes of AAV DJ-CMV eGFP-deleted cre (GVVC-AAV-62) or AAV DJ-CMV eGFP-cre (GVVC-AAV-63) (both generated by the Stanford Gene Vector and Virus Core). The following day, media was fully replaced and 48hrs after infection media was removed and the plate was flash frozen on dry ice and kept in −80 deg. For CRE induction in oligodendrocytes, on day 5 after isolation, SRF-f/f OPCs were plated at a density of 35,000 cells per well in a 12-well plate. On day 6, cells were infected with 5.6-6.6 x 10^8 viral units of a lentivirus-CRE-IRES-GFP or GFP control. On day 7, media was replaced to differentiation media and on day 3 of differentiation the media was removed and plates were frozen at −80.

Cells were scraped in RLT buffer and RNA was extracted with the RNeasy Plus Micro kit (Qiagen, 74034). cDNA and library synthesis were done inhouse using the Smart-seq2 protocol as previously described ^14^ (detailed protocol at https://doi-org.laneproxy.stanford.edu/10.17504/protocols.io.2uvgew6) with several modifications. Due to the low input RNA content, 2μl of RNA extracted from sorted nuclei was reversed transcribed using 18 cycles. Following bead cleanup using 0.7x ratio with AMPure beads (A63881, Fisher), cDNA concentration was measured using the Qubit 1x dsDNA HS kit (Q33231) and normalized to 0.4 ng/μl as input for library prep. 0.4 μl of each normalized sample was mixed with 1.2 μl of Tn5 Tagmentation mix (0.64 μl TAPS-PEG buffer (PEG 8000, V3011, PROMEGA, and TAPS-NaOH pH8.5, BB-2375, Boston Bioproducts), 0.46 μl H2O and 0.1 μl Tn5 enzyme (20034198, Illumina)), then incubated at 55 °C for 10 min. The reaction was stopped by adding 0.4 μl 0.1% sodium dodecyle sulfate (Fisher Scientific, BP166-500). Indexing PCR reactions were performed by adding 0.4 μl of 5 μM i5 indexing primer (IDT), 0.4 μl of 5 μM i7 indexing primer (IDT), and 1.2 μl of KAPA HiFi Non-Hot Start Master Mix (Kapa Biosystems) using 12 amplification cycles. Libraries were purified using two purification rounds with a ratio of 0.8x and 0.7x AMPure beads. Library quantity and quality was assessed using a Bioanalyzer (Agilent) and Qubit. All steps were done manually using 8-strip PCR tubes and PCR reactions were carried out on a 96-plate thermal cycler (Biorad). Libraries were pooled and sequenced on a Nextseq550 sequencer (Illumina) using single end 63bp for Read 1 and 12bp for index 1 with a high output 75bp kit (20024906, Illumina).

Libraries were sequenced to a depth of at least >10 million reads per sample. Raw sequencing files were demultiplexed and known adapters were trimmed with bcl2fastq. Data analysis of raw sequencing data was performed using the nextflow-core RNA-seq pipeline v3.0. Briefly, the core workflow of the pipeline maps filtered reads against the species reference genome using STAR and computes transcript counts using RSEM. For nuclear RNA-seq data, a custom reference genome was created where exon sequences in GTF files were modified to include all introns per transcript and used for the mapping instead. For mouse and rat sequencing data the reference genome GRCm38 and Rnor 6.0 provided by Illumina igenomes were used, respectively. All gene annotations were based on the Ensembl database. Obtained raw gene transcript counts per sample were loaded into DESeq2, performing normalization for transcript length and sequencing depth, and differential expression analysis with standard settings. Effect sizes for each gene were computed based on normalized counts computed by DESeq2 using the function cohen.d of the R package effsize. Gene set enrichment analysis was performed using GeneTrail 3 using BH-FDR p-value adjustment with all remaining parameters kept at default.

### Data Analysis and Statistics

All analysis was conducted blinded to the genotype and experimental condition. Data analysis and statistics were done using GraphPad Prism 9.0 software. Descriptive statistics (mean, standard error of mean, and N) were reported in Figure Legends. Statistical significance was determined between biological replicate (N = 3–5) by an unpaired, two-tailed t-test, unless otherwise stated in the Figure Legends. For in vivo assays, N refers to number of mice. For cellular assays, N refers to number of independent cell purifications from 1-2 mouse brains. We pre-determined that using 3-5 biological replicates for culture or in vivo experiments were sufficient based on previously published studies (Harterink et al., 2017; Zuchero et al., 2015). There were no outliers/excluded data in any experiment.

## Data and materials availability

The data that support the findings of this study and step-by-step protocols are available from the corresponding or first authors upon request. RNAseq and ChIP-seq data will be deposited at appropriate repositories upon publication. Correspondence and requests for all other materials should be addressed to J.B.Z. or T.I.

---

**Supplementary Figure S1.**
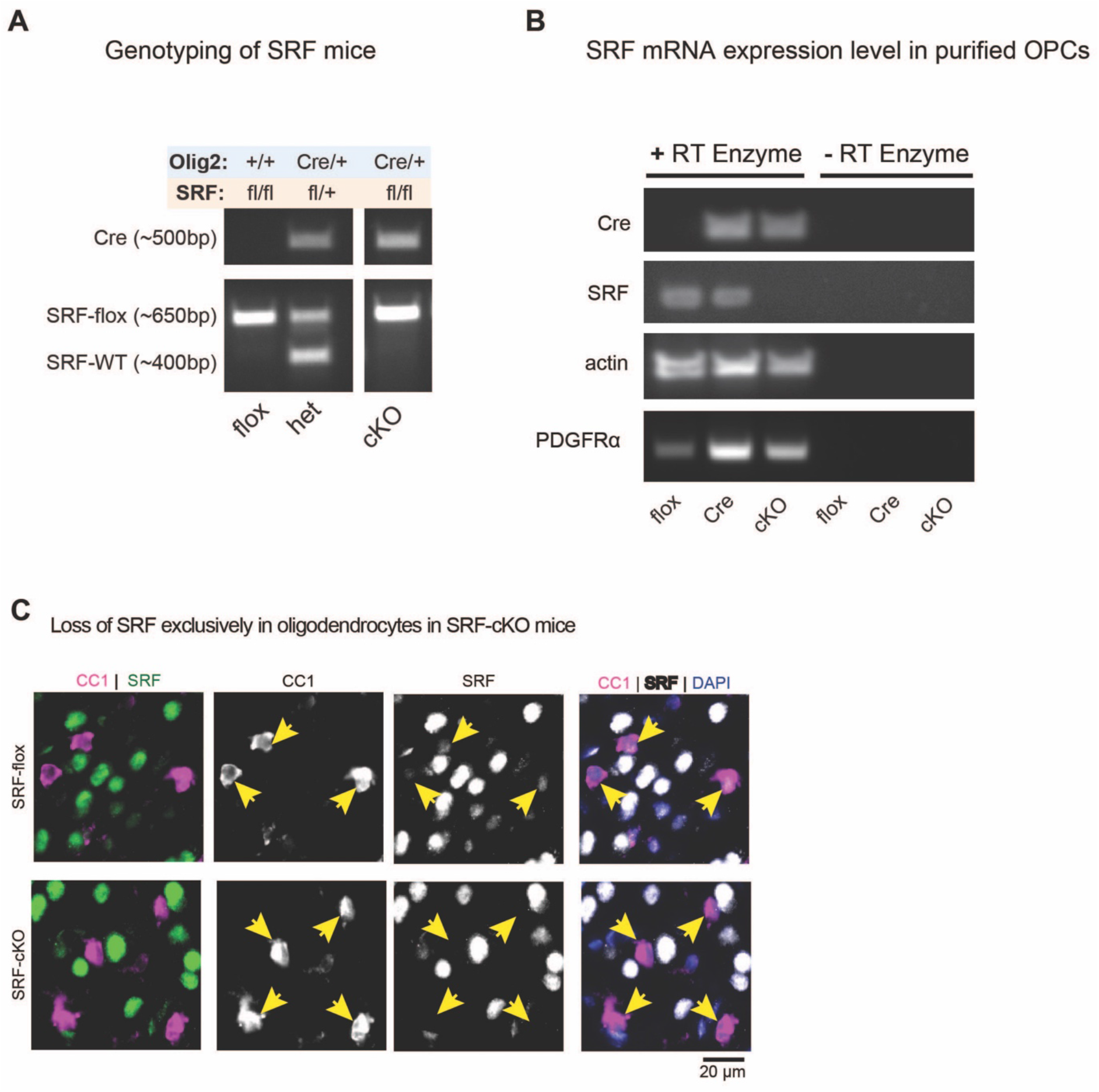
Generation of SRF cKO mice. A. Genotyping strategy and confirmation of Olig2-SRF-cKO mice. B. Validation of loss of Srf from purified OPCs from Srf-flox, CRE mice and SRF-cKO mice by RT-PCR to detect expression of of Cre, SRF, actin and Pdgfra. C. Validation of loss of Srf loss in mature oligodendrocytes (CC1^+^ cells) in brain sections of SRF-flox and SRF-cKO mice by immunostaining.

**Supplementary Figure S2.**
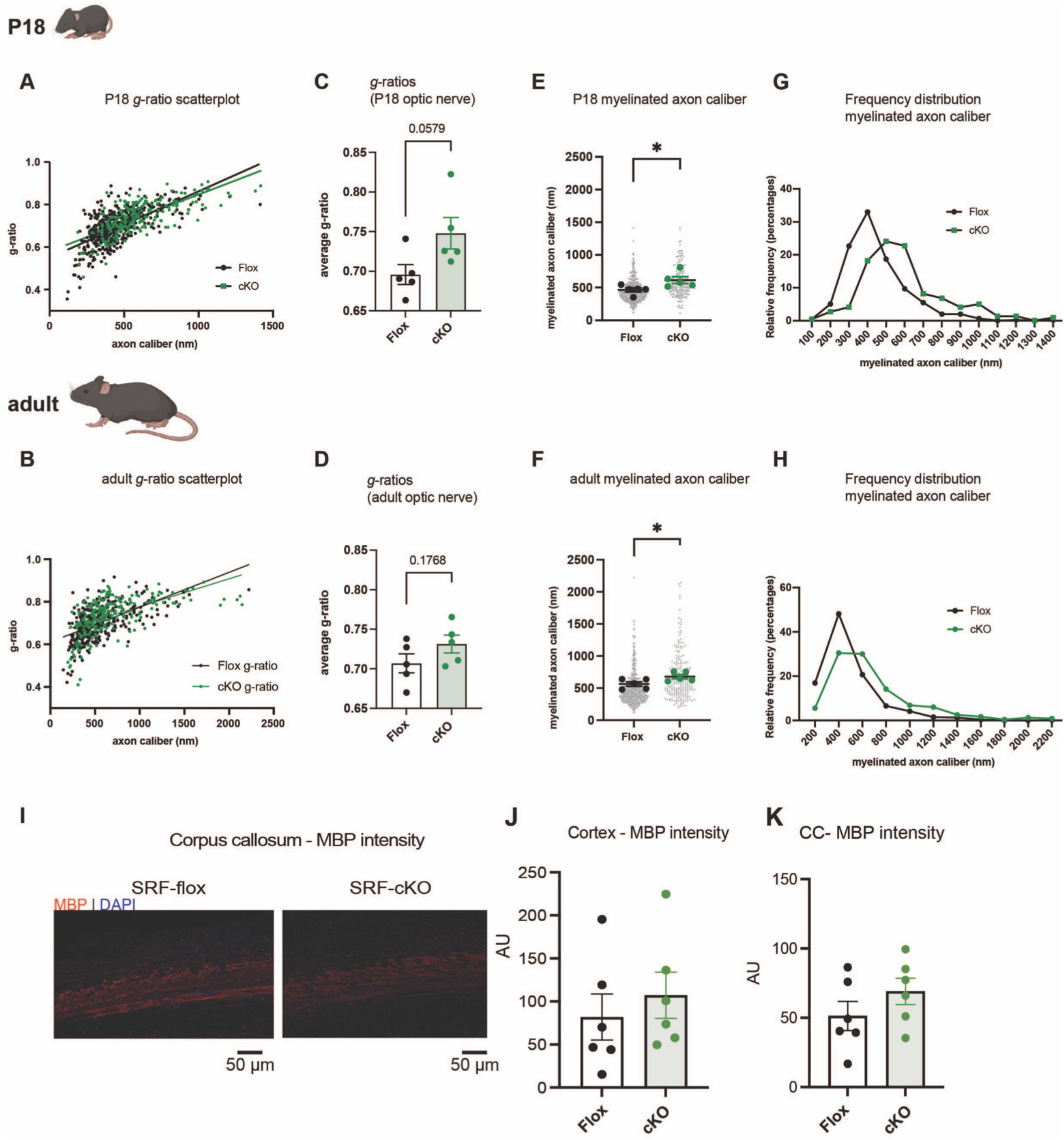
Additional EM analysis of SRF cKO mice. A-B. *g*-ratio of SRF-flox and SRF-cKO optic nerve myelinated axons as a function of axon caliber in P18 (A) and adult (B) mice. C-D. Average *g*-ratios of SRF-flox and SRF-cKO optic nerve myelinated axons at P18 (C) and adult (D). n=5 mice per genotype at each timepoint. E-F. Myelinated axon caliber of SRF-flox and SRF-cKO optic nerve at P18 (E) and adult (F). n=5 mice per genotype at each timepoint. G-H. Frequency distribution of myelinated axon caliber of SRF-flox and SRF-cKO optic nerve axons at P18 (G) and adult (H). I. Representative images of MBP staining in corpus callosum of SRF-flox and SRF-cKO P16-P18 mice. Scale bar represents 50μm. J. Analysis of the mean MBP intensity in the cortex of SRF-flox and SRF-cKO mice. n=6 mice per genotype. Unpaired t-test. K. Analysis of the mean MBP intensity in the corpus callosum (CC) of SRF-flox and SRF-cKO mice. n=6 mice per genotype. Unpaired t-test.

**Supplementary Figure S3.**
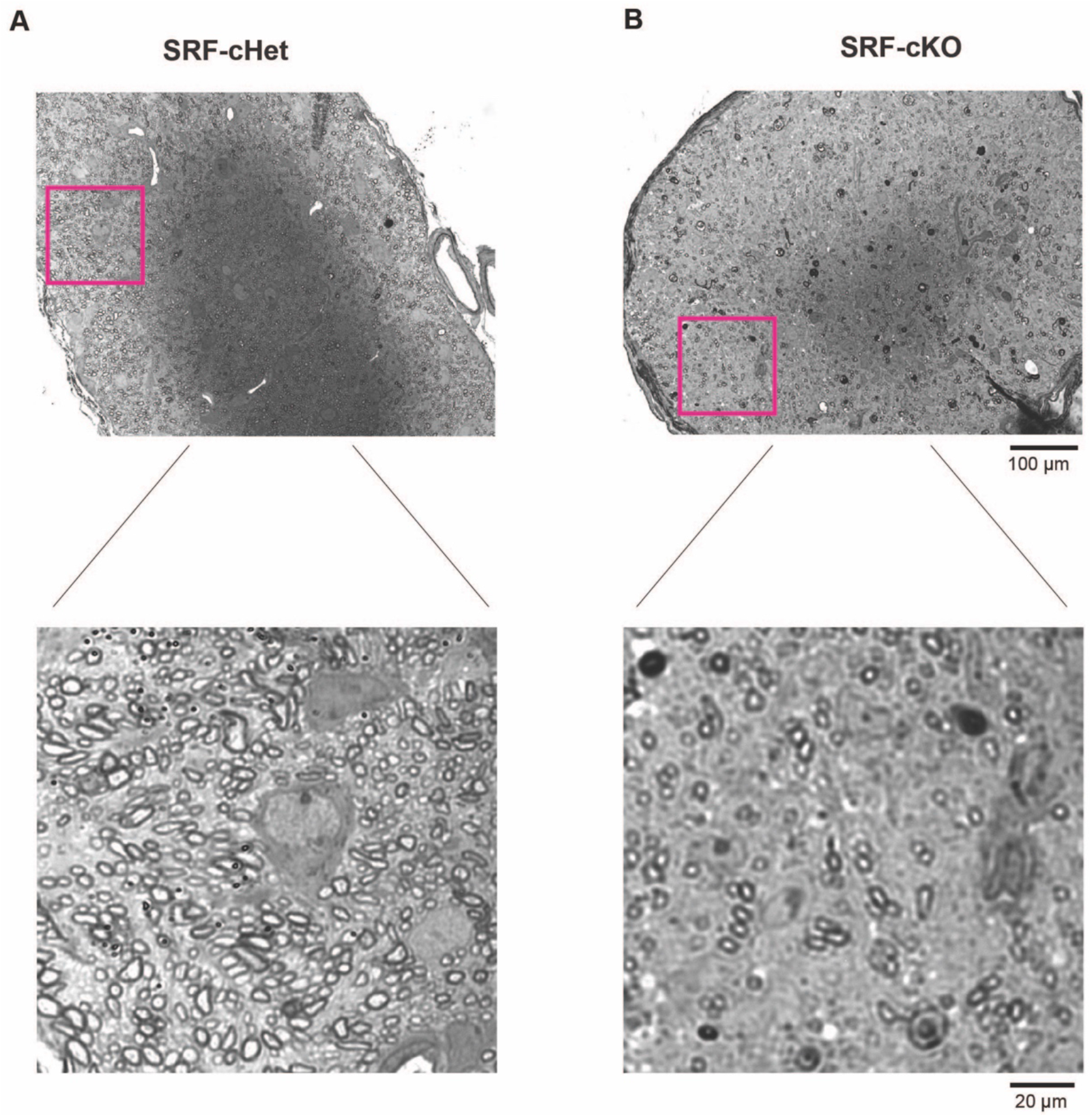
Toluidine blue staining of optic nerve myelin. A-B. Toluidine blue of SRF-cHet (A) and SRF0cKO (B) showing the location of EM analysis in optic nerves of P16 mice. Scale bar represents 100μm and 20μm in insert.

**Supplementary Figure S4.**
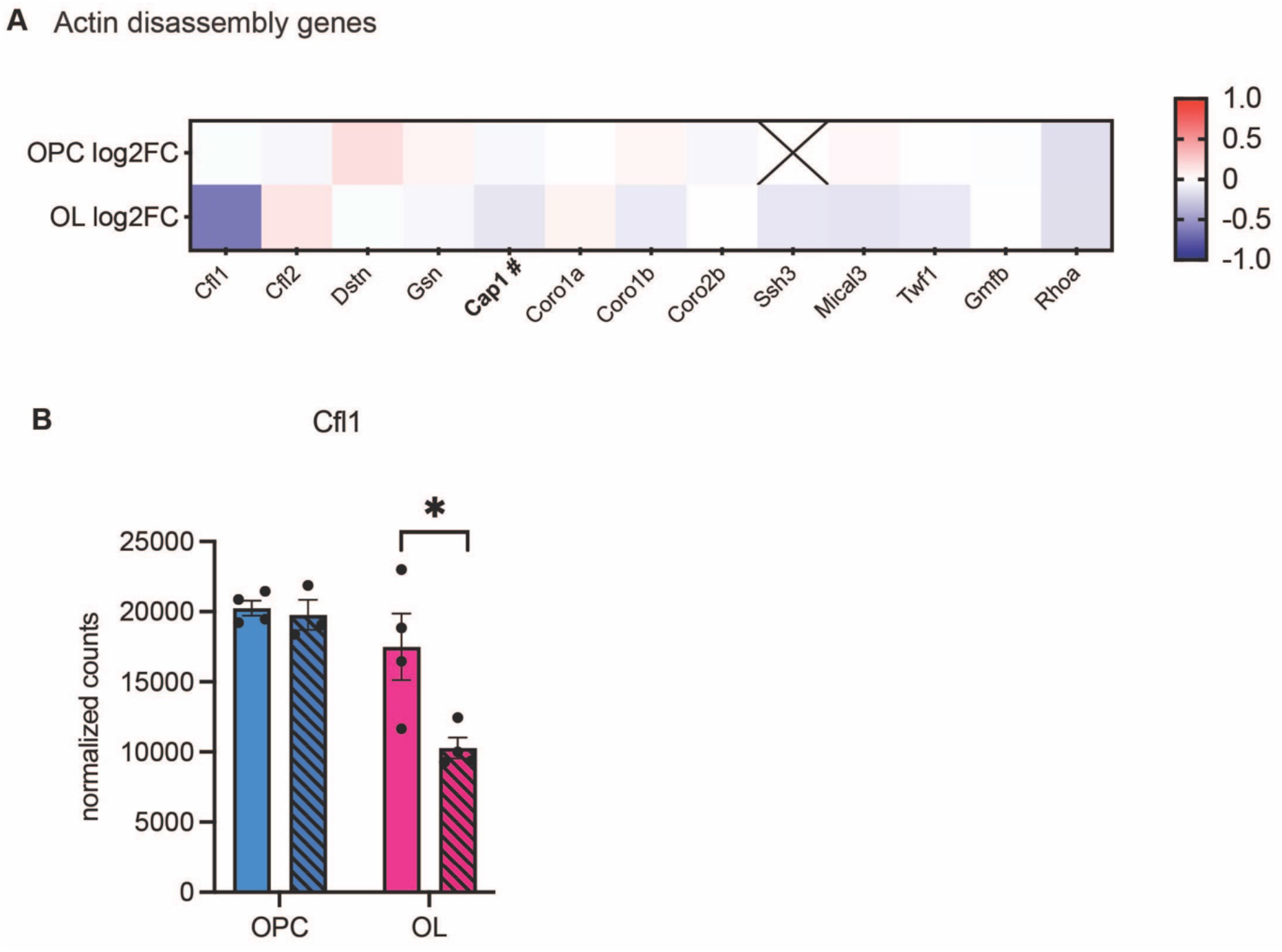
Actin disassembly genes in RNAseq dataset. A. Heatmap of the main actin disassembly genes expressed by OPCs and oligodendrocytes (Zuchero et al., 2015) showing the log2FC of KO vs. flox OPCs and oligodendrocytes. B. Normalized expression counts of Cfl1 in SRF-flox and SRF-KO OPCs and oligodendrocytes. OPC SRF-KO n=3. All the rest n=4. Statistics by Deseq2. # Indicates that Cap1 is a validated hit in the ChIP-seq experiment.

**Supplementary table S1**. Normalized counts and summary statistics of bulk RNAseq of SRF-cKO OPCs vs. SRF-flox OPCs.

**Supplementary table S2.** Normalized counts and summary statistics of bulk RNAseq of SRF-cKO oligodendrocytes vs. SRF-flox oligodendrocytes.

**Supplementary table S3.** SRF ChIP-seq peaks enriched in OPC or oligodendrocyte samples or both (common peaks) including motif analysis details.

## ACKNOWLEDGMENTS

We thank current and past members of the Zuchero, Wyss-Coray, and Barres labs (especially Adiljan Ibrahim, Sophia Wienbar, and Alexandra Münch) and Ben A. Barres for their discussion and support; David Ginty for SRF-floxed mice; David Rowitch for Olig2-Cre mice. Images and diagrams were created using BioRender.com. We gratefully acknowledge the Stanford University Cell Sciences Imaging Core Facility for EM data collection, especially John Perrino and Ibanri Phanwar for processing and staining EM samples (RRID:SCR_017787: supported by ARRA Award Number 1S10RR026780-01 from the National Center for Research Resources [NCRR]. Its contents are solely the responsibility of the authors and do not necessarily represent the official views of the NCRR or the National Institutes of Health). We also thank the Stanford Neuroscience Gene Vector and Virus Core for producing the adeno-associated viruses used in this study.

This project was supported by Wu Tsai Neurosciences Interdisciplinary Scholarships (to TI, MAG, and ML); the Stanford Graduate Fellowship (MI); the Schaller-Nikolich Foundation (A.Ke.); the Michael J. Fox Foundation for Parkinson’s Research (125491594 for A.Ke and F.K.; MJFF-021418 for T.W.-C., A.Ke. and F.K.); the National Institutes of Health AG064897 (TWC), R01NS119823 (JBZ), and NINDS Diversity Supplement R01NS119823-01S1 (MAG); the McKnight Endowment Fund for Neuroscience (JBZ); the National Multiple Sclerosis Society Harry Weaver Neuroscience Scholar Award (JBZ); the Beckman Young Investigator Award (JBZ); the Myra Reinhard Family Foundation (JBZ); the Koret Family Foundation (JBZ); and was originally launched in the lab of Ben A. Barres with support from the National MS Society RG4777-A-10. ML is a Merck-sponsored fellow of the Helen Hay Whitney Foundation.

## AUTHOR CONTRIBUTIONS

TI, MAG, FK, and JBZ designed the study and interpreted the results. TI, MAG, AK, MI, ML, NA, and JBZ performed in vivo and culture experiments and analyzed data. JA, AK, and FK performed and supervised bioinformatics and statistical analysis for RNAseq and ChIP-seq experiments. TI and JBZ wrote the manuscript with feedback from all authors. AK, TWC, FK, and JBZ provided supervision, mentorship, and funding. All authors read and approved the final paper.

## INCLUSION AND DIVERSITY

We worked to ensure sex balance in the selection of non-human subjects. One or more of the authors of this paper self-identifies as an underrepresented ethnic minority in their field of research or within their geographical location. One or more of the authors of this paper received support from a program designed to increase minority representation in their field of research. While citing references scientifically relevant for this work, we also actively worked to promote gender balance in our reference list. We support inclusive, diverse, and equitable conduct of research.

